# Vasodilators activate TMEM16A channels in endothelial cells to reduce blood pressure

**DOI:** 10.1101/2023.06.02.543450

**Authors:** Alejandro Mata-Daboin, Tessa A. C. Garrud, Carlos Fernandez-Pena, Dieniffer Peixoto-Neves, M. Dennis Leo, Angelica K. Bernardelli, Purnima Singh, Kafait U. Malik, Jonathan H. Jaggar

## Abstract

Endothelial cells (ECs) regulate vascular contractility to control regional organ blood flow and systemic blood pressure. Several cation channels are expressed in ECs which regulate arterial contractility. In contrast, the molecular identity and physiological functions of anion channels in ECs is unclear. Here, we generated tamoxifen-inducible, EC-specific *TMEM16A* knockout (*TMEM16A* ecKO) mice to investigate the functional significance of this chloride (Cl^-^) channel in the resistance vasculature. Our data demonstrate that TMEM16A channels generate calcium-activated Cl^-^ currents in ECs of control (*TMEM16A*^*fl/fl*^) mice that are absent in ECs of *TMEM16A* ecKO mice. Acetylcholine (ACh), a muscarinic receptor agonist, and GSK101, a TRPV4 agonist, activate TMEM16A currents in ECs. Single molecule localization microscopy data indicate that surface TMEM16A and TRPV4 clusters locate in very close nanoscale proximity, with ∼18% exhibiting overlap in ECs. ACh stimulates TMEM16A currents by activating Ca^2+^ influx through surface TRPV4 channels without altering the size or density of TMEM16A or TRPV4 surface clusters, their spatial proximity or colocalization. ACh-induced activation of TMEM16A channels in ECs produces hyperpolarization in pressurized arteries. ACh, GSK101 and intraluminal ATP, another vasodilator, all dilate pressurized arteries through TMEM16A channel activation in ECs. Furthermore, EC-specific knockout of TMEM16A channels elevates systemic blood pressure in conscious mice. In summary, these data indicate that vasodilators stimulate TRPV4 channels, leading to Ca^2+^-dependent activation of nearby TMEM16A channels in ECs to produce arterial hyperpolarization, vasodilation and a reduction in blood pressure. We identify TMEM16A as an anion channel present in ECs that regulates arterial contractility and blood pressure.

**One sentence summary:** Vasodilators stimulate TRPV4 channels, leading to calcium-dependent activation of nearby TMEM16A channels in ECs to produce arterial hyperpolarization, vasodilation and a reduction in blood pressure.

## INTRODUCTION

Endothelial cells line the lumen of all blood vessels and regulate several physiological functions, including contractility, which controls regional organ blood flow and systemic pressure. A wide array of receptor ligands, extracellular substances and mechanical stimuli act on endothelial cells to regulate their functions (*1*). Endothelial cells electrically couple to smooth muscle cells in the vascular wall and can directly modulate arterial potential to regulate contractility (*2*). Endothelial cells also produce and release several diffusible vasoactive factors, including nitric oxide (NO), a dilator (*3*). Given that endothelial cells regulate arterial contractility and blood pressure, it is essential to identify novel mechanisms by which vasoactive stimuli modulate their excitability.

Endothelial cells express several cation channels which regulate membrane potential and plasma membrane calcium (Ca^2+^) influx to control arterial contractility. Cation channels present in endothelial cells include TRP vanilloid 4 (TRPV4), TRP polycystin 1 (TRPP1, polycystin-2), TRP ankyrin 1 (TRPA1), and small (SK) and intermediate (IK) conductance Ca^2+^-activated potassium channels (*4-6*). In contrast, the molecular identity and physiological functions of anion channels in endothelial cells are poorly understood. Chloride (Cl^-^) is the predominant extracellular and intracellular anion. Cl^-^ currents were first identified in cultured vascular endothelial cells more than two decades ago (*7-9*). Studies have proposed that Ca^2+^-activated Cl^-^ currents, volume-regulated Cl^-^ currents, and ClC currents occur in cultured endothelial cells (*8, 10-12*). Currently, it is unclear whether non-cultured endothelial cells generate Cl^-^ currents, the molecular identities of the channels that may produce these Cl^-^ currents and their physiological functions.

TMEM16A (anoctamin 1) is a Ca^2+^-activated homodimeric Cl^-^ channel expressed by the *TMEM16A* gene (*13-18*). Recent studies have proposed that TMEM16A channels generate Cl^-^ currents in cultured endothelial cells from tissues including human umbilical vein, heart and brain (*19-21*). Global deletion of *TMEM16A* in mice leads to several phenotypes, including repressed gastrointestinal tract peristalsis and tracheal abnormalities (*22-24*). Constitutive knockout of TMEM16A channels in endothelial cells did not alter the contractility of wire-mounted aortic rings or modify the blood pressure of mice which was measured using a tail-cuff plethysmograph (*19*). Other studies have shown that the constitutive knockout of ion channels can lead to compensatory mechanisms that obscure and complicate attempts to identify their vascular functions (*25-28*). Similarly, the functional significance of TMEM16A channels in the resistance vasculature may differ from that in the aorta, which is a conduit vessel. Thus, alternative approaches to study TMEM16A channels in endothelial cells may reveal that these Cl^-^ channels regulate arterial contractility and blood pressure.

Here, we generated tamoxifen-inducible, endothelial cell-specific TMEM16A knockout mice to investigate the physiological functions of this ion channel in the resistance vasculature. Our data demonstrate that vasodilators activate TRPV4 channels, leading to Ca^2+^ influx which stimulates TMEM16A channels in endothelial cells. TMEM16A channels generate Cl^-^ currents which induce membrane hyperpolarization, vasodilation, and a reduction in blood pressure. We also show that TMEM16A and TRPV4 clusters exhibit nanoscale overlap in the endothelial cell plasma membrane, providing evidence for local Ca^2+^-dependent coupling between these ion channels. These results identify TMEM16A as an anion channel expressed in endothelial cells which regulates arterial contractility and blood pressure.

## RESULTS

### Generation and validation of tamoxifen-inducible, endothelial cell-specific TMEM16A knockout mice

Mice were genetically modified to insert loxP sites between exons 5 and 6 of the *TMEM16A* gene (*TMEM16A*^*fl/fl*^). *TMEM16A*^*fl/fl*^ mice were crossed with *Cdh5*-C*re/ERT2* mice, a tamoxifen-inducible, endothelial cell-specific Cre line, generating *TMEM16A*^*fl/fl*^:ecCre^+^ mice. Genomic PCR indicated that tamoxifen activated *TMEM16A* recombination in *TMEM16A*^*fl/fl*^:ecCre^+^ (*TMEM16A* ecKO) mice but not in *TMEM16A*^*fl/fl*^ mice (Fig. S1A). Western blotting experiments determined that TMEM16A protein in mesenteric arteries of tamoxifen-treated *TMEM16A* ecKO mice was ∼75.8% of that in arteries of tamoxifen-treated *TMEM16A*^*fl/fl*^ mice (Fig. 1A and B). This reduction is expected as TMEM16A is present in other cell types, including arterial smooth muscle cells, where recombination would not occur (*29*). In contrast, TRPV4, endothelial nitric oxide synthase (eNOS), IK, and SK3 proteins were unaltered in arteries of tamoxifen-treated *TMEM16A*^*fl/fl*^:ecCre^+^ mice when compared to those in *TMEM16A*^*fl/fl*^ mice (Fig. 1A and B). TMEM16A immunolabeling was detected in endothelial cells of *en face* mesenteric arteries from tamoxifen-treated *TMEM16A*^*fl/fl*^ mice but negligible in endothelial cells of arteries from *TMEM16A*^*fl/fl*^:ecCre^+^ mice (Fig. 1C).

**Fig. 1.**
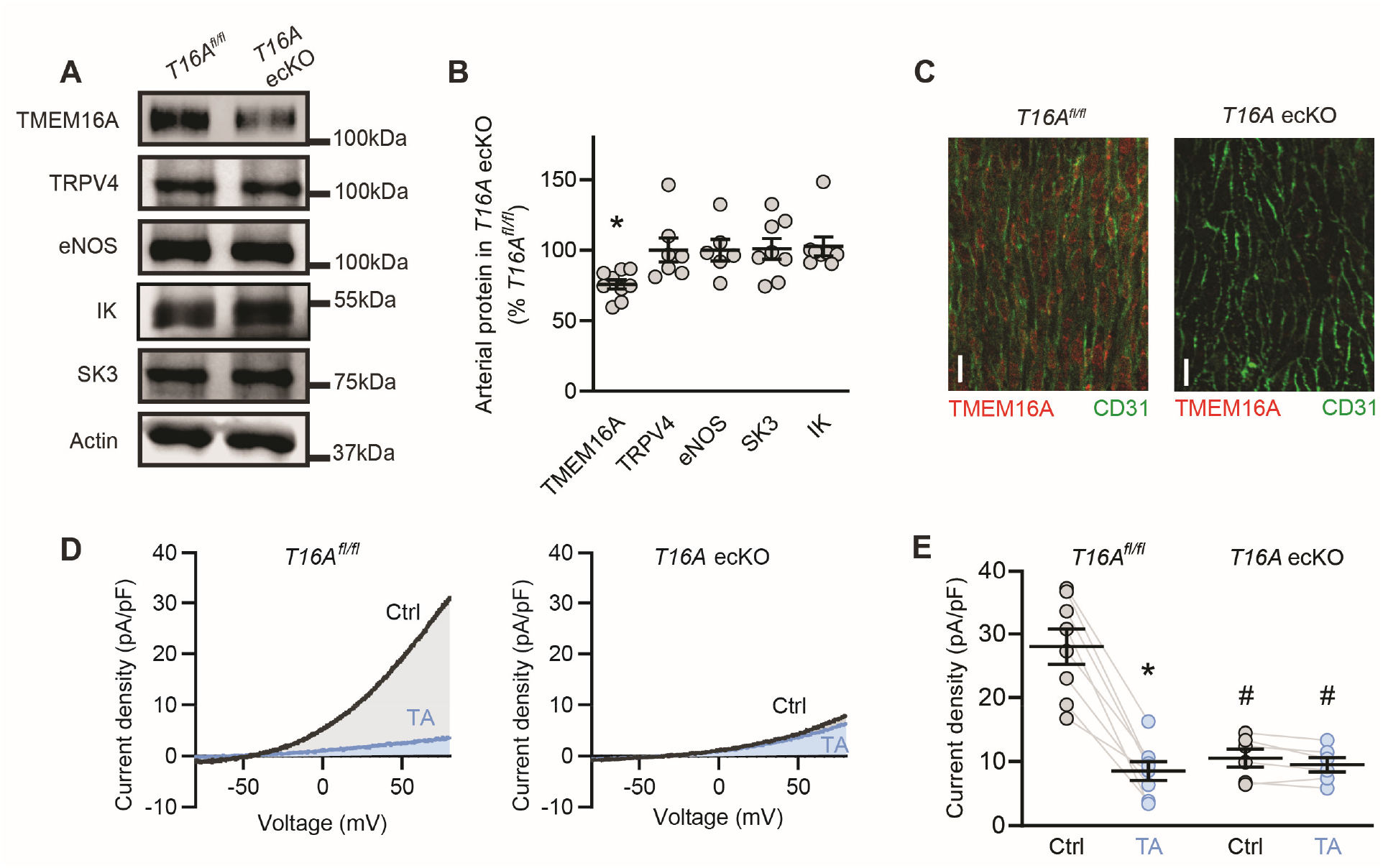
Generation and validation of *TMEM16A* ecKO mice. (A) Western blots illustrating TMEM16A, TRPV4, eNOS, SK3, and IK proteins in mesenteric arteries of tamoxifen-treated *TMEM16A*^*fl/fl*^ (*T16A*^*fl/fl*^) and *TMEM16A* ecKO (*T16A* ecKO) mice. (B) Mean data for TMEM16A (n=9), TRPV4 (n=7), eNOS (n=6), SK3 (n=8), and IK (n=8) proteins in mesenteric arteries of tamoxifen-treated *TMEM16A* ecKO mice when compared to those in tamoxifen-treated *TMEM16A*^*fl/fl*^ mice. * indicates P<0.05 vs *TMEM16A*^*fl/fl*^. (C) *En fac*e immunofluorescence images of endothelial cells from mesenteric arteries in tamoxifen-treated *TMEM16A*^*fl/fl*^ and *TMEM16A* ecKO mice (representative of 4 mesenteric arteries for each genotype). Labelling of CD31, an endothelial cell-specific specific marker, is also shown. Scale bars = 50 µm. (D) Representative whole-cell current recordings of fresh-isolated endothelial cells from *TMEM16A*^*fl/fl*^ and *TMEM16A* ecKO mice with 1 µM free Ca^2+^ in the pipette solution, before and after applying tannic acid (TA, 10 µM) to the bath solution. (E) Mean data obtained from experiments shown in panel D at +80 mV. N=8 for *TMEM16A*^*fl/fl*^. N=6 for *TMEM16A* ecKO. * indicates P<0.05 vs Ctrl within genotype, # indicates P<0.05 in Ctrl *TMEM16A* ecKO vs Ctrl in *TMEM16A*^*fl/fl*^.

### TMEM16A channels produce Ca^2+^-activated Cl^-^ currents in endothelial cells

Patch-clamp electrophysiology was performed to measure whole-cell Cl^-^ currents (I_Cl_) in fresh-isolated mesenteric artery endothelial cells of *TMEM16A*^*fl/fl*^ and *TMEM16A* ecKO mice. With 100 nM free Ca^2+^ present in the pipette solution, I_Cl_ was small in endothelial cells of *TMEM16A*^*fl/fl*^ mice (Fig. S1B and C). Increasing the free Ca^2+^ concentration in the pipette solution from 100 nM to 1 µM elevated I_Cl_ density from 4.1 ± 0.9 to 28.0 ± 2.8 pA/pF, or ∼6.83-fold in *TMEM16A*^*fl/fl*^ endothelial cells (at +80 mV, Fig. S1B and C). Tannic acid, a TMEM16A channel blocker, reduced this Ca^2+^-activated I_Cl_ to ∼30.4 % of control in *TMEM16A*^*fl/fl*^ endothelial cells (at +80 mV, Fig. 1D and E). In contrast, with 1 µM free Ca^2+^ in the pipette solution I_Cl_ was small and insensitive to tannic acid in endothelial cells of *TMEM16A* ecKO mice (Fig. 1D and E). These results indicate that intracellular Ca^2+^ activates TMEM16A channels in endothelial cells of *TMEM16A*^*fl/fl*^ mice but not in endothelial cells of *TMEM16A* ecKO mice.

### ACh activates TRPV4-dependent TMEM16A currents in endothelial cells

Next, we studied the regulation of TMEM16A currents by vasodilators. Whole-cell I_Cl_ was measured using a pipette solution containing 100 nM free [Ca^2+^] and a bath solution containing 2 mM Ca^2+^. Acetylcholine (ACh), an endothelial cell-dependent vasodilator, increased I_Cl_ ∼4.82-fold in endothelial cells of *TMEM16A*^*fl/fl*^ mice (Fig. 2A and B). Tannic acid or benzbromarone, another TMEM16A channel blocker, inhibited the I_Cl_ activated by ACh in *TMEM16A*^*fl/fl*^ endothelial cells (Fig 2A and B, Fig. S2A and B). In contrast, ACh or ACh plus tannic acid did not alter I_Cl_ in *TMEM16A* ecKO endothelial cells (Fig. 2A and B). These data indicate that ACh stimulates TMEM16A-dependent I_Cl_ currents in endothelial cells.

**Fig. 2.**
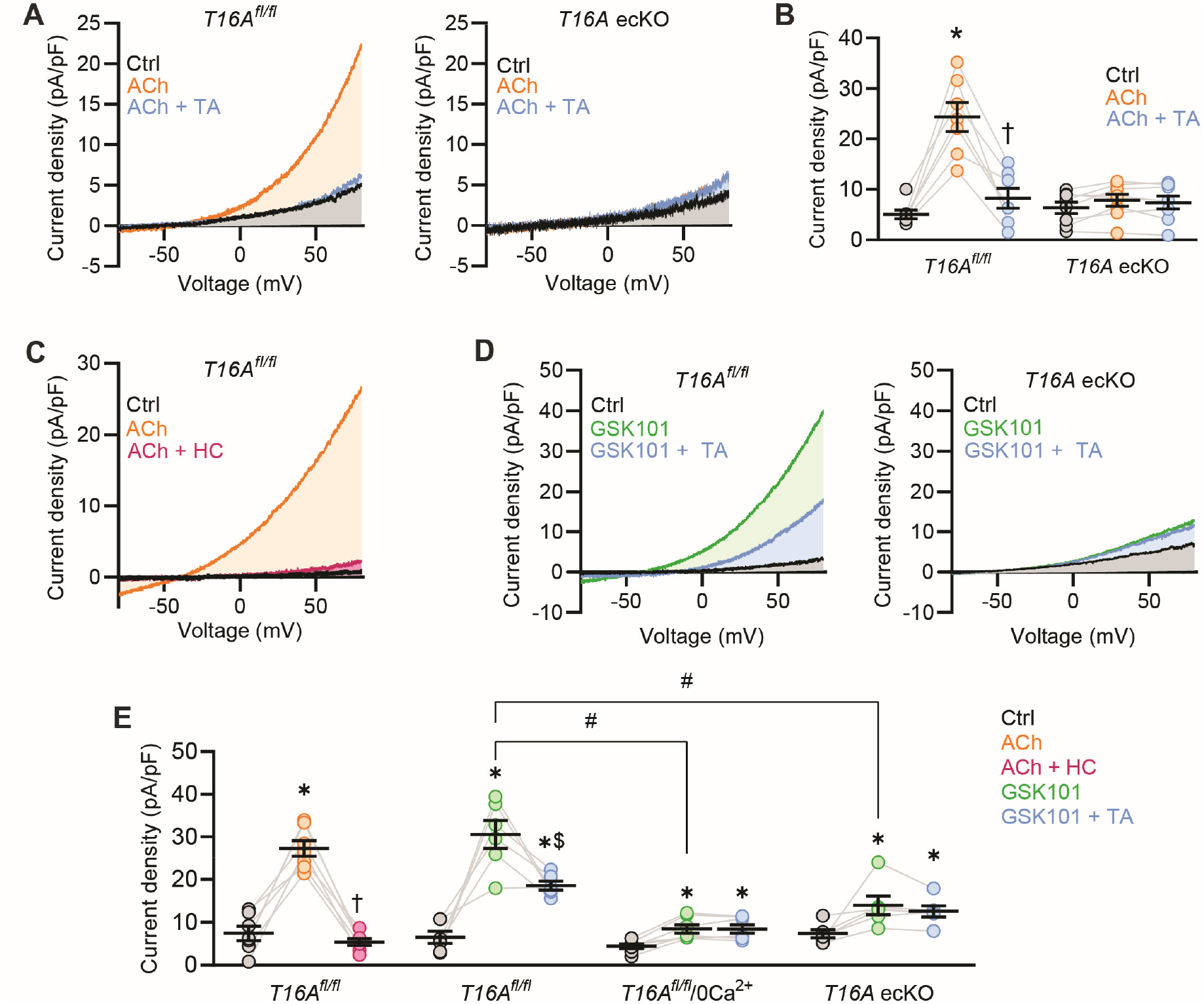
ACh activates TRPV4-dependent TMEM16A currents in ECs. (A) Representative whole-cell current recordings of fresh-isolated endothelial cells from *TMEM16A*^*fl/fl*^ (*T16A*^*fl/fl*^) and *TMEM16A* ecKO (*T16A* ecKO) mice exposed to ACh (10 µM) and ACh + tannic acid (TA, 10 µM). (B) Mean data at +80 mV from experiments in A. N=7 for *TMEM16A*^*fl/fl*^ and n=6 for *TMEM16A* ecKO. * indicates P<0.05 vs Ctrl, † indicates P<0.05 vs ACh (C) Representative whole-cell current recordings of control, ACh (10 µM) and ACh + HC067047 (HC, 1 µM) in the same fresh-isolated endothelial cell from a *TMEM16A*^*fl/fl*^ mouse. (D) Representative whole-cell current recordings of control, GSK101 (1 nM), and GSK101 + TA (10 µM) in the same fresh-isolated endothelial cells from *TMEM16A*^*fl/fl*^ and *TMEM16A* ecKO mice. (E) Mean data at +80 mV from experiments in C (n=7) and D (n=6 for *TMEM16A*^*fl/fl*^ and *TMEM16A* ecKO, n=7 for *TMEM16A*^*fl/fl*^/0 Ca^2+^). * indicates P<0.05 vs Ctrl, † indicates P<0.05 vs ACh, $ indicates P<0.05 vs GSK101, and # indicates P<0.05 vs GSK101 in *TMEM16A*^*fl/fl*^.

ACh activates TRPV4 channels, leading to Ca^2+^ influx in endothelial cells (*6, 30*). We tested the hypothesis that ACh stimulates TRPV4 channels, which activate TMEM16A currents. HC067047, a selective TRPV4 channel blocker, inhibited ACh-induced I_Cl_ activation in endothelial cells of *TMEM16A*^*fl/fl*^ mice (Fig. 2C and E). GSK1016790A (GSK101), a selective TRPV4 channel activator, stimulated a ∼4.70-fold increase in I_Cl_ in *TMEM16A*^*fl/fl*^ endothelial cells (Fig. 2D and E). I_Cl_ activation by GSK101 was similarly inhibited by either tannic acid or the removal of extracellular Ca^2+^ in *TMEM16A*^*fl/fl*^ endothelial cells (Fig. 2D, 2E and Fig. S2C). Inhibition of I_Cl_ by tannic acid or the removal of extracellular Ca^2+^ were also not additive, suggesting a similar mechanism (Fig. S2C, Fig. 2E). In contrast to the effects in *TMEM16A*^*fl/fl*^ cells, GSK101 activated a smaller ∼1.90-fold increase in current that was not altered by tannic acid in *TMEM16A* ecKO endothelial cells (Fig. 2D and E). As cesium (Cs^+^) is the major cation present in the pipette solution, the small GSK101-induced current in *TMEM16A* ecKO endothelial cells may be due to Cs^+^ current through TRPV4 channels (*31*). These results indicate that ACh stimulates TRPV4 channels, leading to Ca^2+^ influx which stimulates TMEM16A channels in endothelial cells.

### TMEM16A and TRPV4 channel surface clusters locate in close spatial proximity in endothelial cells

Single-Molecule Localization Microscopy (SMLM) in combination with Total Internal Reflection Fluorescence (TIRF) imaging was performed to measure the nanoscale properties and spatial proximity of surface TMEM16A and TRPV4 channel clusters in fresh-isolated mesenteric artery endothelial cells. Imaging was performed on cells which labelled for CD31, an endothelial cell-specific marker, in oxygen scavenging photo-switching buffer (Fig. S3A). Localization precision of the Alexa Fluor-conjugated secondary antibodies used to label TMEM16A and TRPV4 channels were calculated to be between 15.9 and 19.1 nm in endothelial cells (Fig. S3B and C). The size of TMEM16A and TRPV4 surface clusters were similar at ∼ 2646 and 2281 nm^2^, respectively, in *TMEM16A*^*fl/fl*^ endothelial cells (Fig. 3A and B). TMEM16A clusters were more numerous than TRPV4 clusters at 20.6 and 13.4 clusters/µm^2^, respectively (Fig. 3C). The mean distance (center to center) between a TMEM16A cluster and its nearest TRPV4 neighbor was ∼107 nm, indicating close spatial proximity (Fig. 3D). Approximately 17.8% of TMEM16A clusters overlapped with a TRPV4 cluster (Fig. 3E). When experimental data were reconstructed using Coste’s randomization algorithm, TMEM16A and TRPV4 overlap was only ∼5.8 % (Fig. 3E). The size and density of TRPV4 clusters were similar in *TMEM16A*^*fl/fl*^ and *TMEM16A* ecKO endothelial cells, indicating that TMEM16A knockout did not alter TRPV4 channel cluster properties (Fig. 3B and C). ACh did not alter the size or density of TMEM16A or TRPV4 clusters, nor their proximity or overlap, indicating that Ca^2+^-dependent signaling between TRPV4 and TMEM16A channels is not associated with changes in the spatial properties of clusters.

**Fig. 3.**
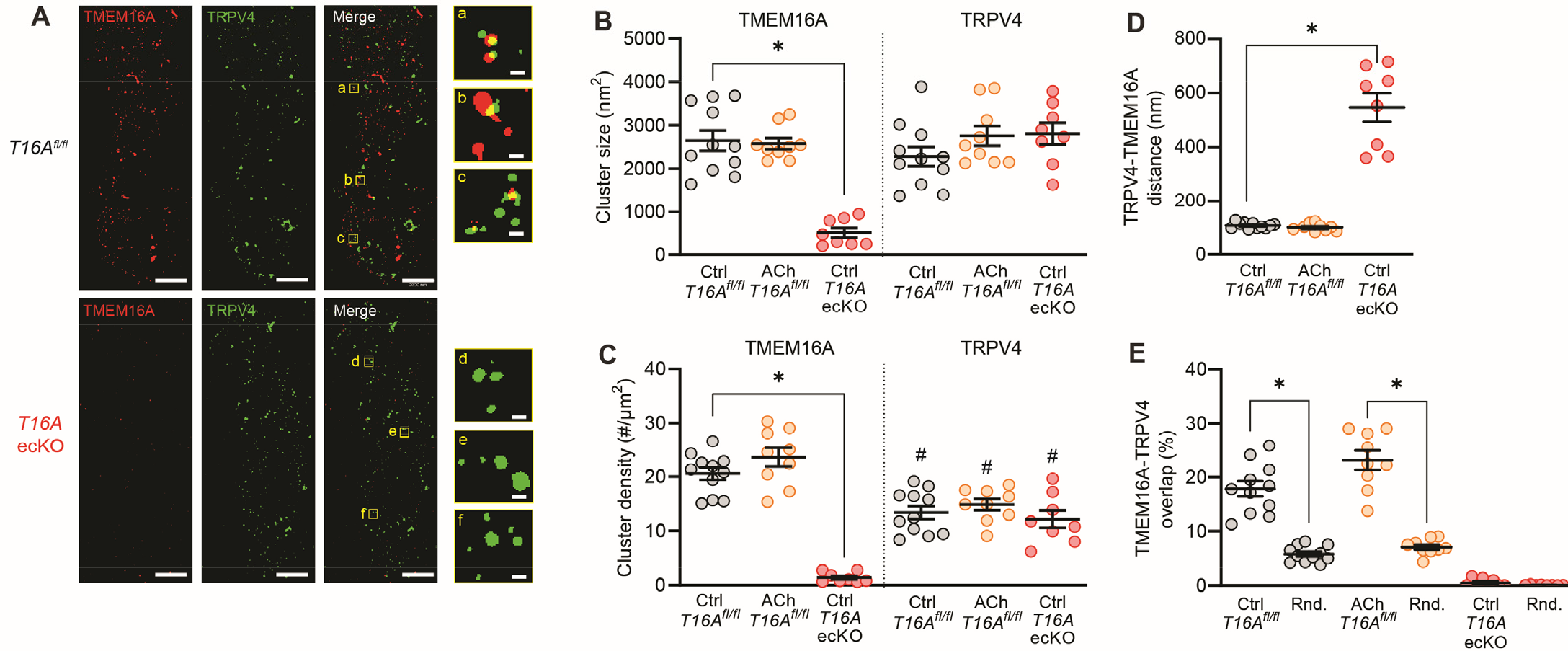
Plasma membrane TMEM16A and TRPV4 clusters locate in nanoscale proximity in ECs. (A) TIRF-SMLM images of TMEM16A and TRPV4 surface clusters in fresh-isolated mesenteric artery endothelial cells from *TMEM16A*^*fl/fl*^ and *TMEM16A* ecKO mice. Scale bars are 2 µm for the whole cell images and 100 nm for the inset images shown on the right. (B) Mean data for TMEM16A and TRPV4 cluster sizes, * indicates P<0.05 vs Ctrl *TMEM16A*^*fl/fl*^. (C) Mean data for TMEM16A and TRPV4 cluster density, * indicates P<0.05 vs Ctrl *TMEM16A*^*fl/fl*^, # indicates P<0.05 vs Ctrl *TMEM16A*^*fl/fl*^. (D) Mean data for TMEM16A to TRPV4 nearest-neighbor cluster analysis, * indicates P<0.05 vs Ctrl *TMEM16A*^*fl/fl*^. (E) Mean data for the percentage of TMEM16A clusters that overlap with a TRPV4 cluster. Also illustrated is TMEM16A to TRPV4 overlap following Coste’s randomization (Rnd) simulation. The number of cells analyzed in each condition is: 11 (*TMEM16A*^*fl/fl*^), 9 (*TMEM16A*^*fl/fl*^ cells exposed to 10 µM ACh) and 8 (*TMEM16A* ecKO), * indicates P<0.05 vs Ctrl for within treatment group.

In *TMEM16A* ecKO endothelial cells, the mean size of TMEM16A clusters was ∼19.0 % of that in *TMEM16A*^*fl/fl*^ endothelial cells (Fig. 3B). TMEM16A clusters were far less numerous in *TMEM16A* ecKO endothelial cells, at ∼6.9 % of that in *TMEM16A*^*fl/fl*^ endothelial cells (Fig. 3C). The distance between TMEM16A and TRPV4 clusters was also much greater at ∼544.8 nm, with only ∼0.5 % of TMEM16A clusters displaying overlap with a TRPV4 cluster in *TMEM16A* ecKO endothelial cells (Fig. 3D and E). These data indicate that clusters observed in *TMEM16A* ecKO cells are due to background labeling of TMEM16A antibodies, validating the data obtained in *TMEM16A*^*fl/fl*^ endothelial cells (Fig. 3A) (*32*). Moreover, these results demonstrate that surface TMEM16A and TRPV4 clusters locate in nanoscale proximity with a significant proportion exhibiting overlap in the endothelial cell plasma membrane.

### Endothelial cell TMEM16A channel activation leads to arterial hyperpolarization and vasodilation

Arterial membrane potential is a major determinant of contractility (*33*). We tested the hypothesis that endothelial cell TMEM16A channels regulate the membrane potential of pressurized (80 mmHg) third-order mesenteric arteries. In control, the membrane potential of *TMEM16A*^*fl/fl*^ and *TMEM16A* ecKO arteries were similar at -32.2 and -31.0 mV, respectively (Fig. 4A and B). ACh stimulated a mean membrane hyperpolarization of ∼ 7.5 mV in *TMEM16A*^*fl/fl*^ arteries (Fig. 4A and B). In contrast, ACh-induced hyperpolarization in *TMEM16A* ecKO arteries was ∼ 2.7 mV, or ∼36.4 % of that in *TMEM16A*^*fl/fl*^ arteries (Fig. 4A and B). Thus, ACh stimulates arterial hyperpolarization, in part, through the activation of endothelial cell TMEM16A channels.

**Fig. 4.**
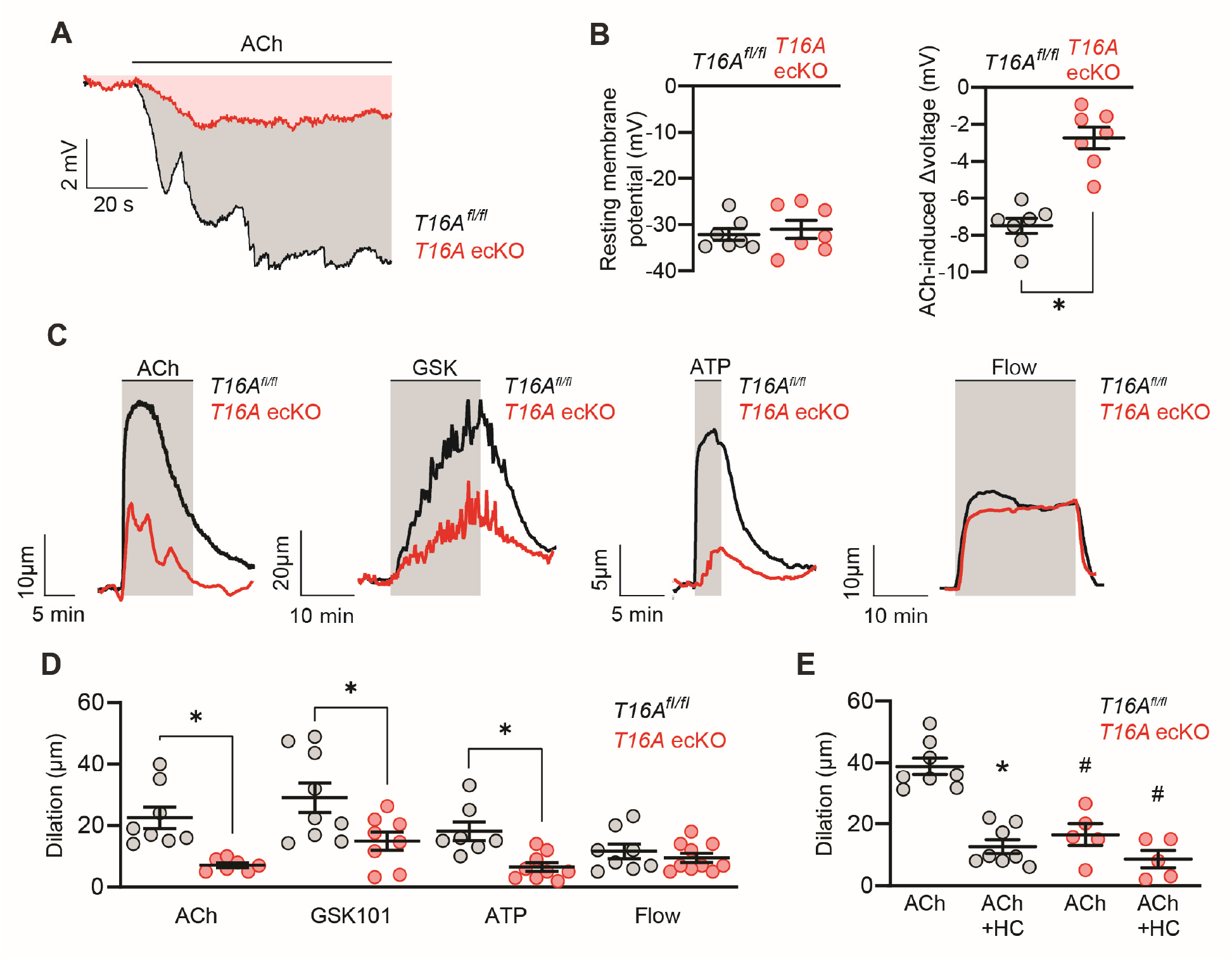
TMEM16A channel activation in endothelial cells contributes to ACh-induced arterial hyperpolarization and vasodilation. (A) Representative membrane potential recordings obtained from microelectrode impalements in pressurized (80 mmHg) mesenteric arteries of *TMEM16A*^*fl/fl*^ and *TMEM16A* ecKO mice showing response to ACh (10 µM). (B) Mean data obtained from experiments in panel A showing the membrane potential in control and the magnitude of the ACh-induced hyperpolarization in arteries of *TMEM16A*^*fl/fl*^ (n=7) and *TMEM16A* ecKO mice (n=7), * indicates P<0.05 vs *TMEM16A*^*fl/fl*^. (C) Representative traces illustrating diameter responses of pressurized (80 mmHg) mesenteric arteries of *TMEM16A*^*fl/fl*^ and *TMEM16A* ecKO mice in response to ACh (10 µM), GSK101 (10 nM), ATP (1 µM) and intravascular flow (15 dyn/cm^2^). (D) Mean data obtained from experiments in panel C showing dilation to ACh (n=8 for *TMEM16A*^*fl/fl*^, n=7 for *TMEM16A* ecKO), GSK101 (n=9 for *TMEM16A*^*fl/fl*^, n=8 for *TMEM16A* ecKO), ATP (n=7 for *TMEM16A*^*fl/fl*^, n=9 for *TMEM16A* ecKO) and intravascular flow (n=8 for *TMEM16A*^*fl/fl*^, n=10 for *TMEM16A* ecKO), * indicates P<0.05 vs *TMEM16A*^*fl/fl*^. (E) Mean data of the inhibitory effect of HC067047 (HC, 1 µM) on ACh-induced vasodilation in pressurized (80 mmHg) mesenteric arteries of *TMEM16A*^*fl/fl*^ (n=8) and *TMEM16A* ecKO mice (n=5), * indicates P<0.05 vs ACh within genotype, # indicates P<0.05 vs ACh *TMEM16A*^*fl/fl*^.

Myography was performed on pressurized (80 mmHg) mesenteric arteries to investigate the regulation of arterial contractility by endothelial cell TMEM16A channels. ACh or intraluminal ATP stimulated vasodilation in *TMEM16A* ecKO arteries that was ∼31.7 and 36.1 %, respectively, of that in *TMEM16A*^*fl/fl*^ arteries (Fig. 4C and D). In contrast, vasodilation to intraluminal flow (15 dyn/cm^2^) was similar in *TMEM16A*^*fl/fl*^ and *TMEM16A* ecKO arteries (Fig. 4C and D). Vasoconstriction to intravascular pressure (myogenic tone), 60 mM K^+^ or phenylephrine, which act on smooth muscle cells, were similar in *TMEM16A*^*fl/fl*^ and *TMEM16A* ecKO arteries (Fig. S4A-D). These data demonstrate that endothelial cell-specific TMEM16A knockout does not cause generalized endothelial cell or smooth muscle cell dysfunction. Consistent with ACh acting via TRPV4 channels to stimulate TMEM16A currents in endothelial cells, GSK101 produced less dilation in *TMEM16A* ecKO mesenteric arteries, at ∼51.6 % of that in arteries of *TMEM16A*^*fl/fl*^ mice (Fig. 4C and D). ACh produced dilation that was substantially inhibited by HC067047 in pressurized arteries of *TMEM16A*^*fl/fl*^ mice and less so in arteries of *TMEM16A* ecKO mice (Fig. 4E). These results indicate that ACh, ATP and TRPV4 channel activation stimulate TMEM16A channels in endothelial cells, leading to membrane hyperpolarization and vasodilation.

### Endothelial cell TMEM16A channels reduce systemic blood pressure

Blood pressure was measured using radiotelemetry in conscious, freely-moving *TMEM16A*^*fl/fl*^ and *TMEM16A* ecKO mice. Prior to tamoxifen-treatment, blood pressure was similar in *TMEM16A*^*fl/fl*^ and *TMEM16A* ecKO mice (Fig. S5). Tamoxifen administration caused a sustained increase in mean arterial pressure in *TMEM16A* ecKO mice but did not alter mean arterial pressure in *TMEM16A*^*fl/fl*^ mice (Fig. 5A). Blood pressure was elevated in *TMEM16A* ecKO mice and occurred during both active (dark) and inactive (light) periods (Fig. 5B). In contrast, heart rate and locomotor activity were similar in *TMEM16A*^*fl/fl*^ and *TMEM16A* ecKO mice (Fig. 5C and D). These results indicate that endothelial cell TMEM16A channels reduce blood pressure in mice.

**Fig. 5.**
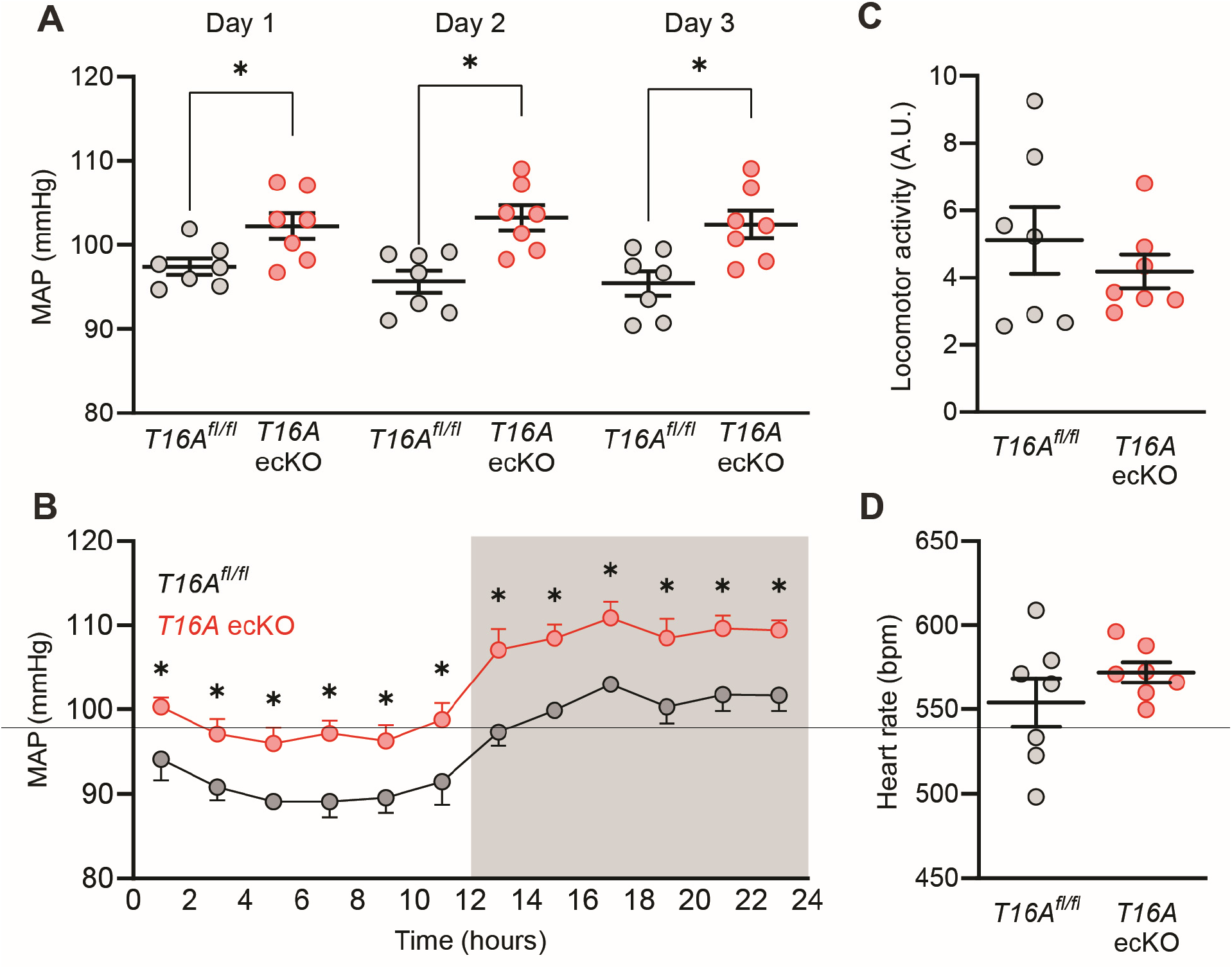
Endothelial cell-specific TMEM16A knockout increases blood pressure. (A) Mean arterial blood pressure (MAP) over a consecutive 3 day time period 14-16 days after the last tamoxifen injection. N=7 per group. * indicates P<0.05 vs *TMEM16A*^*fl/fl*^. (B) MAP measured over a 24 hour period in *TMEM16A*^*fl/fl*^ (n=7) and *TMEM16A* ecKO mice (n=7) 14 days after the last tamoxifen injection. * indicates P<0.05 versus *TMEM16A*^*fl/fl*^. (C) Mean data of locomotor activity. N=7 per group. (D) Mean heart rate. N=7 per group.

## DISCUSSION

Here, we generated a tamoxifen-inducible, endothelial cell-specific TMEM16A knockout mouse model to investigate the physiological functions of this Cl^-^ channel in this cell type. We show that ACh and ATP activate TMEM16A channels in endothelial cells, leading to vasorelaxation. TMEM16A and TRPV4 protein clusters locate in nanoscale proximity in the plasma membrane of endothelial cells, with a significant number exhibiting overlap. ACh activates TRPV4 channels, leading to Ca^2+^ influx which stimulates TMEM16A channels. TMEM16A channel activation in endothelial cells produces membrane hyperpolarization and vasodilation. We also demonstrate that TMEM16A channel knockout in endothelial cells elevates blood pressure in mice.

Cl^-^ currents have been recorded in cultured endothelial cells, including those from the aorta and pulmonary arteries (*34*). Ca^2+^-activated Cl^-^ currents, volume-regulated Cl^-^ currents, and ClC currents occur in cultured endothelial cells (*8, 10-12*). Cultured neonatal mouse cardiac endothelial cells, cultured brain endothelial cells and cultured human umbilical vein endothelial cells generate Cl^-^ currents which are sensitive to manipulations that include TMEM16A inhibitors, TMEM16A-targeted RNAi, and a TMEM16A pore-blocking antibody (*19-21*). Whether non-cultured endothelial cells generate Cl^-^ currents, the molecular identity of the proteins that generate these currents and their physiological functions were unclear. Here, using genetic knockout and pharmacological approaches we show that TMEM16A channels generate Ca^2+^-activated Cl^-^ currents in fresh-isolated endothelial cells.

ACh activates TRPV4 channels in endothelial cells, resulting in Ca^2+^ influx which stimulates SK and IK channels to induce vasodilation (*6*). Our data show that TRPV4-mediated Ca^2+^ influx activates TMEM16A channels in endothelial cells. We did not determine the mechanism by which ATP activates TMEM16A channels in mesenteric artery endothelial cells to cause vasodilation, but purinergic receptors activate TRPV4 channels in pulmonary artery endothelial cells, providing one explanation for our data (*35*). Our data suggest that endothelial cell TMEM16A channels do not contribute to flow-mediated vasodilation. Mechanical stimulation using a fluid stream was shown to activate a Cl^-^ current in cultured human aortic endothelial cells, although the molecular identity of the channels involved was not determined (*36*). Recent studies have demonstrated that flow stimulates a polycystin-1/polycystin-2 (TRPP1) complex in endothelial cells, leading to eNOS, SK, and IK channel activation and vasodilation (*4, 32*). Flow also stimulates Piezo1, a Ca^2+^-permeant non-selective cation channel, in both arterial and capillary endothelial cells (*37, 38*). Our data therefore suggest that PC-1/PC-2 and Piezo1 do not signal to TMEM16A in endothelial cells. Flow may also activate Cl^-^ currents through different signaling mechanisms in non-cultured mesenteric artery endothelial cells and cultured aortic endothelial cells, indicating that further investigation of the mechanisms involved is required.

The binding of ACh to Gα_q/11_-coupled muscarinic receptors can stimulate both TRPV4 channels on the plasma membrane and IP_3_ receptors on the endoplasmic reticulum, leading to an increase in intracellular Ca^2+^ concentration (*6*). We show that inhibition of TRPV4 channels blocks ACh-induced TMEM16A currents, pharmacological activation of TRPV4 channels using GSK101 activates TMEM16A currents, and removing extracellular Ca^2+^ inhibits TRPV4-mediated TMEM16A channel activation. These data indicate that Ca^2+^ influx through TRPV4 channels stimulates TMEM16A currents in endothelial cells. We did not investigate whether IP_3_ receptor-mediated Ca^2+^ release also activates TMEM16A currents. Such an investigation is more suitable for a future study, particularly as the relative contributions of these Ca^2+^ signaling mechanisms may depend on several factors, including the agonist used, its concentration, and the cell voltage, which determines the driving force for plasma membrane Ca^2+^ influx.

SMLM imaging demonstrated that ∼18 % of TMEM16A clusters overlap with a TRPV4 cluster, with the average distance from a TMEM16A cluster to its nearest TRPV4 neighbor 107 nm. At physiological voltages, the EC_50_ for Ca^2+^ of TMEM16A channels is in the low micromolar concentration range (*14*). In contrast, global intracellular Ca^2+^ concentration ranges between 34 and 220 nM nanomolar in non-cultured ECs (*39*). Thus, TMEM16A channels are likely to be activated by local intracellular Ca^2+^ transients generated by nearby TRPV4 channels, which are termed sparklets (*6*). Our data support this concept as the pipette solution in our patch-clamp experiments contained EGTA, a slow Ca^2+^ chelator that buffers global Ca^2+^ signals but does not abolish local Ca^2+^ transients produced by Ca^2+^ channels (*40*). It is possible that TMEM16A channels are activated by Ca^2+^ signals generated by other plasma membrane ion channels and by organellar Ca^2+^ channels, such as IP_3_ receptors. Ca^2+^ influx through TRPV4 channels has also been reported to trigger Ca^2+^-induced Ca^2+^ release from IP_3_Rs (*41*). Such Ca^2+^-induced Ca^2+^ release may activate TMEM16A channels which are more distant from TRPV4 channels.

ACh produced membrane hyperpolarization in pressurized arteries of *TMEM16A*^*fl/fl*^ mice that was attenuated in arteries of *TMEM16A* ecKO mice. For TMEM16A channel activation to elicit hyperpolarization, the equilibrium potential for Cl^-^ (E_Cl_) must be negative of physiological voltage to create a driving force for Cl^-^ influx. Intracellular Cl^-^ concentration in non-cultured endothelial cells of pressurized mesenteric arteries does not appear to have been previously measured but it can be estimated when considering arterial membrane potential. At an intravascular pressure of 80 mmHg and with 122 mM Cl^-^ in the bath solution, the membrane potential of *TMEM16A*^*fl/fl*^ pressurized mesenteric arteries was ∼-32.2 mV. Thus, intracellular Cl^-^ must be less than 34.8 mM for TMEM16A currents to produce hyperpolarization in endothelial cells.

TRPV4 channel activation stimulates SK and IK channels and eNOS signaling in endothelial cells, leading to membrane hyperpolarization and vasodilation (*6, 28, 30*). We now add TMEM16A channels as a functional Ca^2+^-activated downstream target of TRPV4 channels in endothelial cells. Endothelial cell-specific TMEM16A channel knockout reduced ACh- and ATP-induced vasodilation by two-thirds. SK and IK channels are also major targets of TRPV4 channels in endothelial cells (*6*). These observations raise the question of how the loss of one TRPV4 channel target can dramatically reduce agonist-induced vasodilation. One explanation is that TRPV4-mediated SK, IK, and TMEM16A channels may act both in series and in parallel to amplify functional responses. Membrane hyperpolarization caused by SK, IK and TMEM16A activation would increase the driving force for TRPV4-mediated Ca^2+^ influx, further stimulating each channel type. In agreement with this concept, IK and SK channels act in a positive-feedback manner to amplify ACh-induced Ca^2+^ signaling in endothelial cells, with the loss of one K^+^ channel type sufficient to inhibit ACh-induced Ca^2+^ signaling (*42*). Here, our results show that genetic ablation of TMEM16A in endothelial cells robustly attenuates the magnitude of ACh-induced membrane hyperpolarization and vasodilation. This may occur through the inhibition of a positive-feedback mechanism which attenuates SK and IK channel activation, and perhaps eNOS activation. Future studies should investigate these possibilities.

Our data, obtained with a tamoxifen-inducible model of endothelial cell-specific TMEM16A knockout, are in contrast to those acquired with a constitutive (non-inducible) TMEM16A knockout mouse model (*19*). Constitutive TMEM16A knockout did not alter ACh-induced relaxation in wire-mounted, pre-constricted aortic rings (*19*). Similarly, blood pressure in constitutive TMEM16A knockout mice was the same as in control mice when measured using tail-cuff plethysmography (*19*). Constitutive TMEM16A knockout may have produced signaling compensation in mice that nullified a contractility and blood pressure phenotype. Similar examples include mice with constitutive knockout of TRPC6, TRPM4 and TRPV4 channels, which produced paradoxical effects on vascular contractility and blood pressure associated with compensatory mechanisms (*25-28*). We studied resistance-size arteries that develop myogenic tone and regulate systemic blood pressure. In contrast, the effects of constitutive TMEM16A knockout was examined in the aorta, a conduit vessel that does not develop myogenic tone or blood pressure and a vessel in which endothelial cell TMEM16A channels may not regulate contractility. Preconstriction of wire-mounted aortic rings may have also inhibited TMEM16A channels in endothelial cells, thereby preventing their subsequent regulation of contractility. We also show that inducible TMEM16A knockout increased blood pressure which was recorded using telemetry. Tail cuff plethysmography is generally considered to be a less sensitive method than radiotelemetry to measure blood pressure, which may also explain the lack of a phenotype in the constitutive TMEM16A channel knockout (*43, 44*).

A recent study screened several commonly used TMEM16A channel inhibitors (CaCCinh-A01, MONNA, Ani9, niclosamide, and niflumic acid) for selectivity (*45*). The authors suggested that all but one inhibitor, Ani9, blocked the sarco/endoplasmic reticulum (SR/ER) Ca^2+^ ATPase and IP_3_ receptors, which interfered with Ca^2+^ signaling (*45*). Here, we used benzbromarone and tannic acid, which were not screened in the aforementioned study. We also compared the effects of these inhibitors on both *TMEM16A*^*fl/fl*^ and *TMEM16A* ecKO endothelial cells, which provides an important control. We conclude that Ca^2+^ influx through TRPV4 stimulates TMEM16A channels and did not examine the regulation of TMEM16A channels by ER Ca^2+^ release.

In summary, we show that physiological and pharmacological vasodilators activate TMEM16A channels in endothelial cells. Surface TMEM16A clusters locate in nanoscale proximity with TRPV4 clusters and exhibit overlap, and Ca^2+^ influx through TRPV4 channels stimulates TMEM16A channels to elicit Cl^-^ currents. We also show that TMEM16A channel activation in endothelial cells causes arterial hyperpolarization, vasodilation, and a reduction in blood pressure

## MATERIALS AND METHODS

### Animals

All procedures were performed in accordance with the Institutional Animal Care and Use Committee (IACUC) at the University of Tennessee Health Science Center. Mice with loxP sites inserted between exons 5 and 6 of the *TMEM16A gene* (*TMEM16A*^*fl/fl*^) were custom-made for us by Taconic-Cyagen. *TMEM16A*^*fl/fl*^ mice were crossed with *Cdh5-Cre/ERT2* mice, a tamoxifen-inducible, EC-specific Cre line, generating *TMEM16A*^*fl/fl*^:ecCre+ mice. *Cdh5(PAC)-CreERT2* mice were kindly provided by Cancer Research UK (*46*). The genotypes of all mice were confirmed using PCR before use (Transnetyx). All mice (male, 12 weeks of age) were injected with tamoxifen (50 mg/kg, i.p.) once per day for 5 days and studied between 14 and 21 days after the last injection.

### Tissue preparation and EC isolation

Mice were anesthetized with isoflurane (1.5%) followed by decapitation. First to fifth order mesenteric arteries were harvested, cleaned of adventitial tissue and then immersed in ice-cold physiological solution with the following concentrations (in mM): 134 NaCl, 6 KCl, 2 CaCl_2_, 1 MgCl_2_, 10 HEPES, 10 glucose (pH 7.4). Endothelial cells were isolated by enzymatic dissociation of mesenteric arteries by incubating in a Ca^2+^-free solution containing (in mM): 80 Na glutamate, 55 NaCl, 5.6 KCl, 10 HEPES, 10 glucose, 2 MgCl_2_, pH 7.4 with 2 mg/ml protease and 0.33 mg/ml hyaluronidase for 20 minutes at 37 °C. Elastase was included for the last 5 minutes of incubation at a concentration of 0.1 mg/ml. After multiple washes using ice-cold Ca^2+^-free isolation solution, arteries were minced and gently triturated. Isolated endothelial cells were maintained in ice-cold isolation solution and used within 6 hours.

### Genomic PCR

Genomic DNA was isolated from brain homogenate using a Purelink Genomic DNA kit (Thermo Fisher Scientific). Reaction conditions used were initial denaturation at 95 °C for 2 minutes followed by 45 cycles at 95 °C for 0.5 min, 55 °C for 0.5 min, 72 °C for 5 min, and extension at 72 °C for 10 min. The primer sequences used to identify floxed and deleted alleles, in tamoxifen-treated TMEM16A^fl/fl^ or *TMEM16A* ecKO mice, were: Forward 5’-CTCAGGCAATCTCAGTGAAGC-3’ and Reverse 5’-GAACTGTCCTGGAGACACAGG-3’.

### Western blotting

Whole mesenteric arteries were dissected and placed into RIPA buffer (Sigma-Aldrich: R0278) containing protease inhibitor cocktail (Sigma-Aldritch P8340, 1:100 dilution). Arteries were cut with scissors then homogenized (Argos technologies A0001 tissue homogenizer). The resulting lysate was centrifuged at 4 °C, 12,000 rpm for 10 minutes, supernatant was then collected and assayed for protein concentration. Samples were heated in Laemelli Buffer (Biorad 1610747) at 100°C for 5 minutes prior to gel loading. Samples were run on 7.5% SDS-polyacrylamide gels then proteins were transferred onto nitrocellulose membranes. Membranes were blocked with 5% milk for 1 hour then incubated overnight at 4 °C with primary antibody. Membranes were washed and incubated with secondary antibodies at room temperature for 45 minutes. ECL substrate (Thermofisher 34095) was added to membranes and bands were imaged using a ChemiDoc Touch Imaging System (Bio-Rad). Quantification of protein bands was done using ImageJ (NIH) software. All protein bands were normalized to the corresponding actin band.

### En-face arterial immunofluorescence

Arteries were cut longitudinally and fixed with 4% paraformaldehyde in PBS for 1 hour. After washing with PBS, the arteries were permeabilized with 0.2% TritonX-100, blocked with 5% goat serum and incubated overnight at 4 °C with anti-TMEM16A (Cell Signaling, Rabbit D1M9Q 1:100) and CD31 (Abcam, Rat ab7388 1:100) primary monoclonal antibodies. Arteries were incubated with Alexa fluor 488 donkey anti-rat secondary antibody (1:400; Thermo Fisher) and Alexa Fluor 555 donkey anti-rabbit secondary antibody (1:400; Thermo Fisher) for 1 hour at room temperature. Segments were washed with PBS, oriented on slides with the endothelial layer downwards and mounted in 80% glycerol solution. Alexa 488 and Alexa 555 were excited at 488 and 555 with emission collected at 488-540 nm and ≥555 nm, respectively, using a Zeiss LSM 710 laser-scanning confocal microscope.

### Patch-clamp electrophysiology

Fresh-isolated endothelial cells were allowed to adhere to a glass coverslip in a recording chamber. Macroscopic Cl^-^ currents were measured using the conventional whole-cell configuration applying voltage ramps (0.13 mV/ms) between -80 to +80 mV from a holding potential of -40 mV. The pipette solution contained (in mM): 10 CsCl, 130 Cs aspartate, 10 HEPES, 10 glucose, 1 EGTA, 1 MgATP, and 0.2 NaGTP (pH 7.2 adjusted with CsOH). Total Mg^2+^ was adjusted to give a final free concentration of 1 mM while total Ca^2+^ was adjusted to give a final free concentration of 100 nM or 1 uM, depending on the experimental condition. Free Mg^2+^ and Ca^2+^ were calculated using WebmaxC Standard (https://somapp.ucdmc.ucdavis.edu/pharmacology/bers/maxchelator/webmaxc/webmaxcS.htm). The bath solution contained 140 NMDG-Cl, 10 HEPES, 10 glucose, 1 MgCl_2_, and 2 CaCl_2_ (pH 7.4). Ca^2+^-free bath solution was the same composition as the bath solution except Ca^2+^ was omitted and 1 mM EGTA added. Membrane currents were recorded using an Axopatch 200B amplifier, Digidata 1332A, and Clampex 10.3 (Molecular Devices). Currents were filtered at 5 kHz and normalized to membrane capacitance. Liquid junction potential was corrected offline. All experiments were performed at room temperature (22°C). Offline analysis was performed using Clampfit 10.4.

### Single-molecule localization microscopy

Fresh-isolated mesenteric artery endothelial cells we allowed to adhere to 35 mm glass-bottom dishes were fixed in 4% paraformaldehyde and permeabilized with 0.5% Triton X-100 PBS solution. After blocking with 5 % BSA, the cells were inmunolabelled overnight at 4 °C using primary antibodies to TMEM16A, CD31 and TRPV4. Secondary antibodies, Alexa Fluor 488 donkey anti-mouse, Alexa Fluor 555 goat anti-rat, and Alexa Fluor 647 donkey anti-rabbit, were used for detection. ONI Bcubed buffers A+B (ONI) were mixed at the time of experiment, added to the cells to acquire the images, and replaced every hour. A super-resolution Zeiss Elyra 7 microscope equipped with 488 nm (500 mW), 561 nm (500 mW), and 642 nm (500 mW) lasers was used to capture the images with a 63x Plan-Apochromat (NA 1.46) oil immersion lens and a CMOS camera. The camera was operated in frame-transfer mode at a rate of 100 Hz with an exposure time of 30 ms. TIRF mode was used with emission band-pass filters of 550– 650 and 660–760 nm.

CD31 labeling (Alexa 488) was acquired with lattice-SIM to identify endothelial cells that were then used for SMLM imaging. Reconstruction of lattice-SIM images was performed using the SIM processing tool of ZEN software (Zeiss black edition).

Localization precision of fluorophores was calculated using σ^2^_x,y_ =[(s^2^+q^2^/12)/N]+[(8πs^4^b^2^)/(q^2^N^2^)], where σ_x,y_ is the localization precision of a fluorescent probe in lateral dimensions, s is the standard deviation of the point-spread function, N is the total number of photons gathered, q is the pixel size in the image space, and b is the background noise per pixel. The precision of localization is proportional to DLR/√N, where DLR is the diffraction-limited resolution of a fluorophore and N is the average number of detected photons per switching event, assuming the point-spread functions are Gaussian. 30,000 images were used to reconstruct TMEM16A and TRPV4 clusters. This was done by fitting signals to a Gaussian function and applying a point spread function calculated with a standardized 40 nm bead slide using ZEN Black software (Zeiss). The first 2500 frames were excluded to allow for the photo-switching probes to stabilize. Software drift correction was applied using model-based cross-correlation. Cluster properties and co-localization were analyzed using ImageJ v2.1.0 (NIH) and JaCoP v2.0 plugin.

### Pressurized artery myography

Isolated third-order mesenteric arteries segments, 1-2 mm in length, were cannulated at each end in a perfusion chamber (Living Systems Instrumentation) continuously perfused at 37 °C with physiological saline solution (PSS) that contained (in mM): 112 NaCl, 6 KCl, 24 NaHCO_3_, 1.8 CaCl_2_, 1.2 MgSO_4_, 1.2 KH_2_PO_4_ and 10 glucose, gassed with 21% O_2_, 5% CO_2_ and 74% N_2_ to maintain pH at 7.4. A Servo pump model PS-200-P (Living Systems) was used to modify intravascular pressure, which was monitored through pressure transducers. Arterial diameter was recorded at a frequency of 1 Hz using a Nikon TS100-F microscope equipped with a CCD camera and the automatic edge-detection function of IonWizard software (Ionoptix). Myogenic tone was calculated as: 100×(1−Dactive/Dpassive) where Dactive is active arterial diameter and Dpassive is the diameter determined in the presence of Ca^2+^-free PSS supplemented with 5 mM EGTA.

### Pressurized artery membrane potential measurements

Membrane potential was measured in pressurized (80 mmHg) myogenic third-order mesenteric arteries following the development of stable myogenic tone. A sharp glass microelectrode with a resistance between 50-90 MOhm was filled with 3M KCl and inserted into the adventitial side of the arterial wall. Successful intracellular impalements were determined by the following criteria: a sharp negative deflection in potential upon insertion, voltage stability for at least one minute after entry, a sharp positive voltage deflection upon exit from the recorded cell and a < 10% change in tip resistance after the impalement. Membrane potential was acquired using a WPI FD223a amplifier and digitized using a MiniDigi 1A USB interface controlled by pClamp 9.2 software (Axon Instruments).

### Telemetric blood pressure and locomotion measurements

Radiotransmitters (PA-C10, Data Sciences International) were implanted subcutaneously into anesthetized mice, with the sensing electrode placed in the aorta via the left carotid artery. Mice were allowed to recover for 7-10 days. Blood pressure was then recorded every 10 s for 7 days prior to tamoxifen injection and again for 14 days after the last tamoxifen injection (50 mg/kg per day for 5 consecutive days, i.p) using a PhysioTel Digital telemetry platform (Data Sciences International). Dataquest A.R.T. software was used to acquire and analyze data.

### Statistical analysis

Statistical analysis was performed using Graphpad Prism software. Data are expressed as means ± SE. Paired and unpaired data from two populations were compared using Student t-test, while ANOVA with Tukey post hoc test was used for multiple group comparisons. Significance was considered at P<0.05. When P>0.05 a power analysis was applied to verify that the sample size gave a value >0.8.

## Supplementary Materials

Fig. S1. Validation of *TMEM16A*^*fl/fl*^ and *TMEM16A* ecKO mice.

Fig. S2. TMEM16A current properties in ECs.

Fig. S3. Identification of ECs for SMLM and localization precision of the fluorescent indicators used in endothelial cells.

Fig. S4. Smooth muscle-specific vasoconstriction is unaltered in *TMEM16A*^*fl/fl*^ ecKO arteries.

Fig. S5. Mean arterial pressure before tamoxifen injections is similar in *TMEM16A*^*fl/fl*^ and *TMEM16A* ecKO mice.

## Acknowledgments

We thank the Advanced Imaging Core at the University of Tennessee Health Science Center for technical assistance during super resolution imaging experiments.

## Author Contributions

Conceptualization: JHJ, AM-D

Methodology: AM-D, TACG, CF-P, DP-N, MDL, AKB, PS

Investigation: AM-D, TACG, CF-P, DP-N, MDL, AKB, PS

Funding acquisition: JHJ, KUM, AM-D, TACG

Project administration: JHJ

Supervision: JHJ, KUM

Writing – original draft: JHJ, AM-D, TACG

Writing – review & editing: JHJ, KUM, AM-D, TACG, CF-P, DP-N, MDL, AKB, PS

## Data and Materials Availability

All data are available in the main text or the supplementary materials.

