## Supplement for "Vasodilators activate TMEM16A channels in endothelial cells to reduce blood pressure"

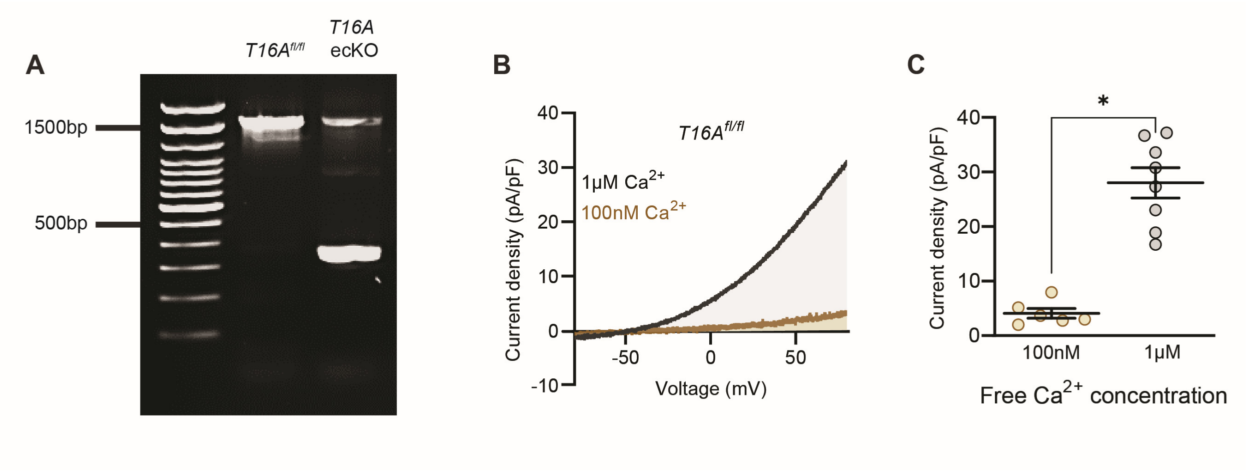


**Fig. S1. Validation of *TMEM16A$^{fl/fl}$* and *TMEM16A* ecKO mice.** (A) Ethidium bromide gel illustrating PCR products of brain homogenates in tamoxifen-treated *TMEM16A$^{fl/fl}$* and *TMEM16A* ecKO mice. Tamoxifen-treated *TMEM16A$^{fl/fl}$* mice produced a ~1,500 bp transcript containing the recombination site. Genomic DNA of tamoxifen-treated *TMEM16A* ecKO mice amplified two transcripts of ~1,500 bp and ~350 bp. The smaller PCR product reflects TMEM16A recombination in endothelial cells, whereas the larger PCR transcript is amplified from TMEM16A present in other cell types, where recombination would not occur. (B) Representative whole-cell current recordings from fresh-isolated endothelial cells of *TMEM16A$^{fl/fl}$* mice with 100 nM free $Ca^{2+}$ (brown) or 1 µM free $Ca^{2+}$ (black) in the pipette solution. (C) Mean data from experiments shown in panel A measured at +80 mV (* indicates $P<0.05$ vs100 nM).

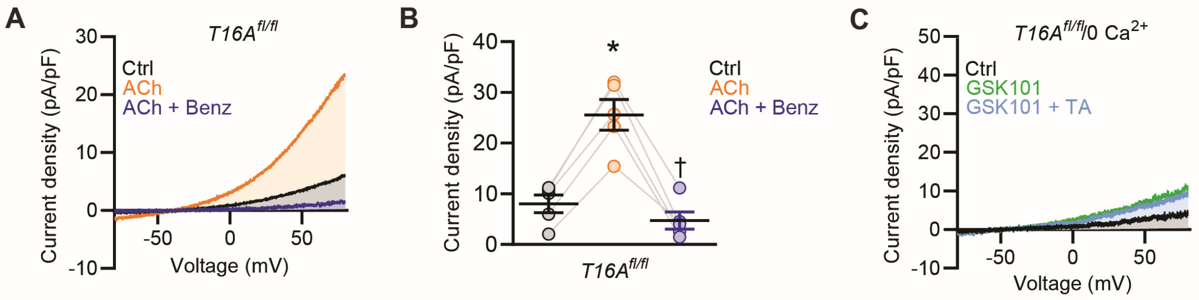


**Fig. S2. TMEM16A current properties in ECs.** (A) Representative whole-cell current recordings of fresh-isolated endothelial cells from *TMEM16A*$^{fl/fl}$ mice exposed in control (Ctrl), ACh (10 µM) and ACh + benzbromarone (Benz, 5 µM). (B) Mean data obtained from experiments shown in panel A measured at +80 mV (n=5). * indicates P<0.05 vs control (Ctrl), † indicates P<0.05 versus ACh. (C) Representative recordings illustrating the regulation of whole-cell current by GSK101 (1 nM) and GSK101 + TA (10 µM) in the absence of extracellular $Ca^{2+}$ in fresh-isolated endothelial cells of *TMEM16A*$^{fl/fl}$ mice.

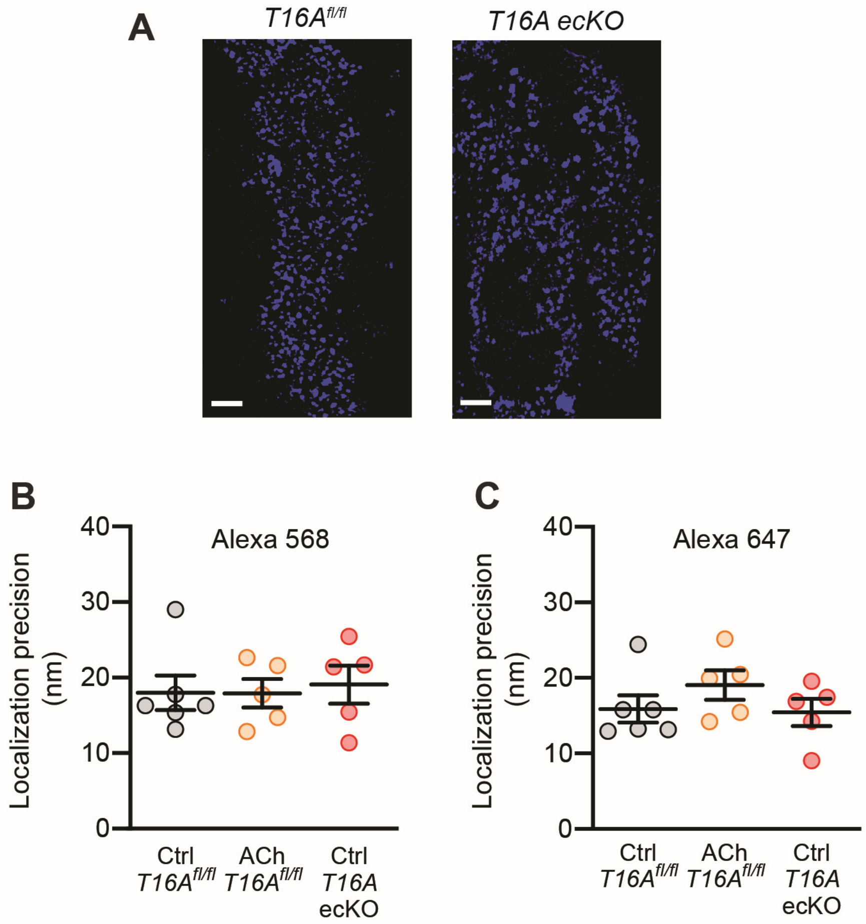


**Fig. S3. Identification of ECs for SMLM and localization precision of the fluorescent indicators used in endothelial cells.** (A) Lattice-SIM images illustrating fluorescence emitted by CD31 coupled to Alexa 488 in cells used for SMLM. Scale bars = 2 µm. (B) Mean data illustrating localization precision of Alexa Fluor 568 in endothelial cells of TMEM16A$^{fl/fl}$ mice in control (Ctrl) and ACh and in cells of *TMEM16A* ecKO mice in control (Ctrl). (C) Mean data illustrating localization precision of Alexa Fluor 647 in the indicated experimental conditions. N=6 in *TMEM16A$^{fl/fl}$*, n=5 in *TMEM16A$^{fl/fl}$* + ACh and n=5 in *TMEM16A* ecKO.

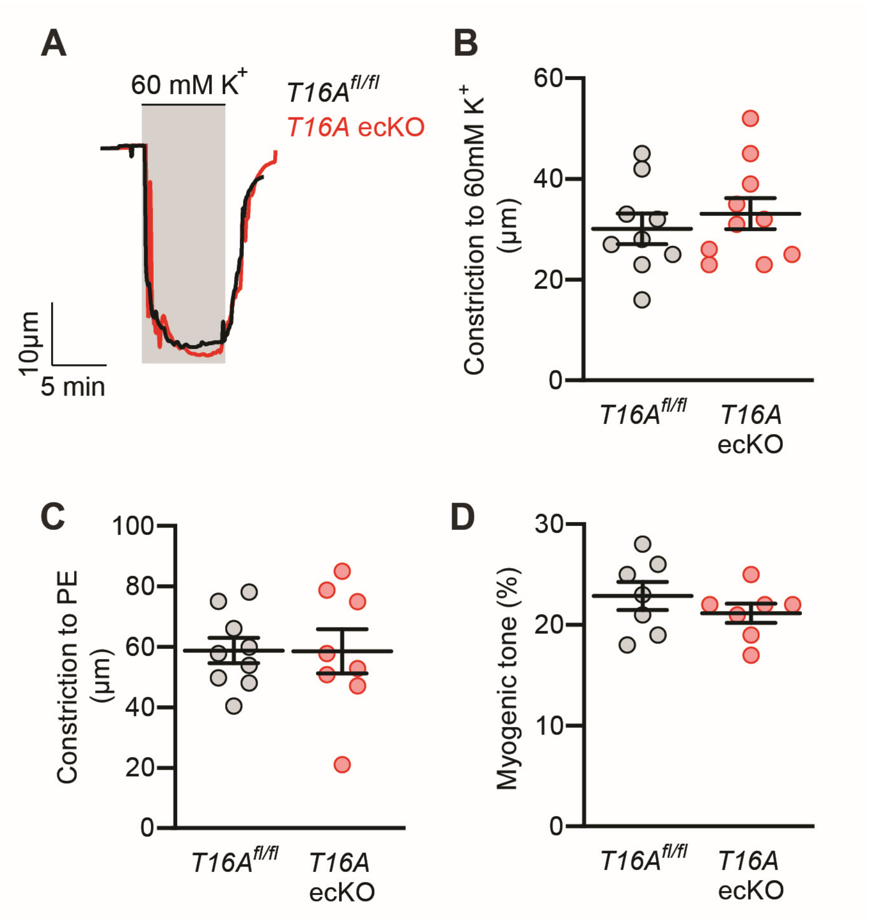


**Fig. S4. Smooth muscle-specific vasoconstriction is unaltered in *TMEM16A*$^{fl/fl}$ ecKO arteries.** (A) Representative traces illustrating constriction to 60 mM $K^+$ in pressurized (10 mmHg) mesenteric arteries of *TMEM16A*$^{fl/fl}$ and *TMEM16A* ecKO mice. (B) Mean data obtained from panel A experiments. N=9 for *TMEM16A*$^{fl/fl}$. N=10 for *TMEM16A* ecKO. (C) Mean data illustrating constriction to PE (phenylephrine, 1 µM). N=9 for *TMEM16A*$^{fl/fl}$. N=8 for *TMEM16A* ecKO. (D) Mean myogenic tone in pressurized (80 mmHg) mesenteric arteries from *TMEM16A*$^{fl/fl}$ and *TMEM16A* ecKO mice. N=7 per group.

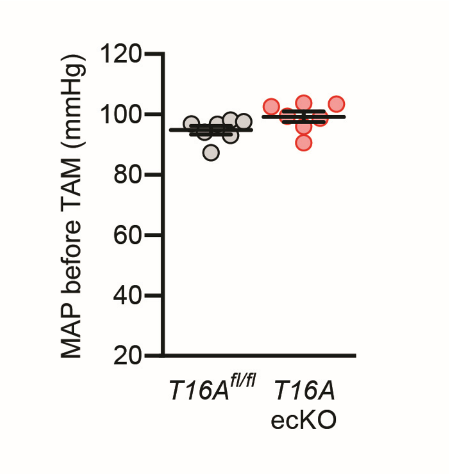


**Fig. S5. Mean arterial pressure before tamoxifen injections is similar in *TMEM16A$^{fl/fl}$* and *TMEM16A* ecKO mice.** Mean arterial pressure data in *TMEM16A$^{fl/fl}$* and *TMEM16A* ecKO mice before tamoxifen injections. N=7 per group.
